# The dot-compartment revealed? Diffusion MRI with ultra-strong gradients and spherical tensor encoding in the living human brain

**DOI:** 10.1101/584730

**Authors:** Chantal M.W. Tax, Filip Szczepankiewicz, Markus Nilsson, Derek K. Jones

**Affiliations:** Cardiff University Brain Research Imaging Centre (CUBRIC), School of Psychology, Cardiff University, Cardiff, UK; Radiology, Brigham and Women’s Hospital, Boston, MA, USA; Harvard Medical School, Boston, MA, USA; Medical Radiation Physics, Clinical Sciences Lund, Lund University, Lund, Sweden; Radiology, Clinical Sciences Lund, Lund University, Lund, Sweden; Mary MacKillop Institute for Health Research, Australian Catholic University, Melbourne, Australia

## Abstract

The so-called “dot-compartment” is conjectured in diffusion MRI to represent small spherical spaces, such as cell bodies, in which the diffusion is restricted in all directions. Previous investigations inferred its existence from data acquired with directional diffusion encoding which does not permit a straightforward separation of signals from ‘sticks’ (axons) and signals from ‘dots’. Here we combine isotropic diffusion encoding with ultra-strong diffusion gradients (240 mT/m) to achieve high diffusion-weightings with high signal to noise ratio, while suppressing signal arising from anisotropic water compartments with significant mobility along at least one axis (e.g., axons). A dot-compartment, defined to have apparent diffusion coefficient equal to zero and no exchange, would result in a non-decaying signal at very high b-values (b ≳ 7000 s/mm^2^). With this unique experimental setup, a residual yet slowly decaying, signal above the noise floor for b-values as high as 15000 s/mm^2^ was seen clearly in the cerebellar grey matter (GM), and in several white matter (WM) regions to some extent. Upper limits of the dot-signal-fraction were estimated to be ~2% in cerebellar GM and ~0.2% in WM. By relaxing the assumption of zero diffusivity, the signal at high b-values in cerebellar GM could be represented more accurately by an isotropic water pool with a low apparent diffusivity of 0.11 µm^2^/ms and a substantial signal fraction of ~7-16%. This remaining signal at high b-values has potential to serve as a novel and simple marker for isotropically-restricted water compartments in cerebellar GM.

## 1. Introduction

Diffusion Magnetic Resonance Imaging (dMRI) (Le Bihan and Breton, 1985) probes structures at much smaller length-scales than the imaging resolution by sensitising the signal to the random molecular motion of water. Biophysical modelling of the contributions to this signal aims to characterise tissue microstructure properties by carefully selecting model compartments (typically multiple non-exchanging water pools) that have a measurable impact on the signal (Stanisz et al., 1997). In healthy white matter (WM), biophysical models typically include anisotropic extra- and intra-axonal compartments (Alexander et al., 2010; Assaf and Basser, 2005; Fieremans et al., 2011; Jespersen et al., 2007; Kroenke et al., 2004; Lampinen et al., 2019; Novikov et al., 2018; Sotiropoulos et al., 2012; Stanisz et al., 1997; Zhang et al., 2012). The inclusion of a so-called “dot-compartment” for WM-modelling is motivated by the observation of an almost constant, non-attenuating signal at very high b-values (e.g., b ≳ 7000 s/mm^2^). This has been hypothesised to arise from the ubiquity of small isotropic spaces (e.g., glial cell-bodies) wherein the diffusion of water molecules is highly restricted in all directions (Alexander et al., 2010; Stanisz et al., 1997), leading to a near-zero apparent diffusivity. A method to measure the signal fraction of such isotropically-restricted components accurately *in vivo* could thus potentially provide a proxy for the density of cells and enable quantification of cellular pathology in a wide range of neurological and psychiatric disorders.

Previous work investigating compartmental contributions to the dMRI signal from conventional pulsed-gradient encoding – also called Stejskal-Tanner encoding (Stejskal and Tanner, 1965) or linear tensor encoding (LTE (Westin et al., 2016)) – showed that including a dot-compartment provided a more complete description of the WM dMRI signal, both *ex vivo* (Panagiotaki et al., 2012) and *in vivo* (Ferizi et al., 2014; Zeng et al., 2018). However, a dot-compartment is not generally included in WM biophysical models, e.g. (Assaf and Basser, 2005; Behrens et al., 2003; Jespersen et al., 2007; Kroenke et al., 2004; Novikov et al., 2018; Zhang et al., 2012). Moreover, a recent study of the dMRI signal in WM at b-values up to 10000 s/mm^2^ on a clinical MRI system suggested that the WM dot-signal-fraction is negligible (Veraart et al., 2019).

Probing the dot-compartment in anisotropic tissue is challenging with conventional LTE, due to the strong relationship between encoding-direction and orientation-distribution of anisotropic tissue microenvironments. Even when measuring along the dominant axis of a fibre bundle in which there is orientation dispersion, a slow diffusing component can be observed (due to the gradient direction not being perfectly parallel to all of the fibres); it is therefore challenging to disentangle this from the scenario in which a dot-compartment is present (Fig.1a). Here, we address this problem by the use of spherical tensor encoding (STE, also called isotropic diffusion encoding) to render signals insensitive to orientation and anisotropy (Eriksson et al., 2013; Lasič et al., 2014; Mori and Van Zijl, 1995; Westin et al., 2016; Wong et al., 1995). STE at high b-values can suppress the dMRI signal from water pools that are mobile along at least one axis (Fig.1a). At sufficiently high b-values only the signal from compartments with extremely low or zero diffusivity in all directions would remain.

**Fig. 1:**
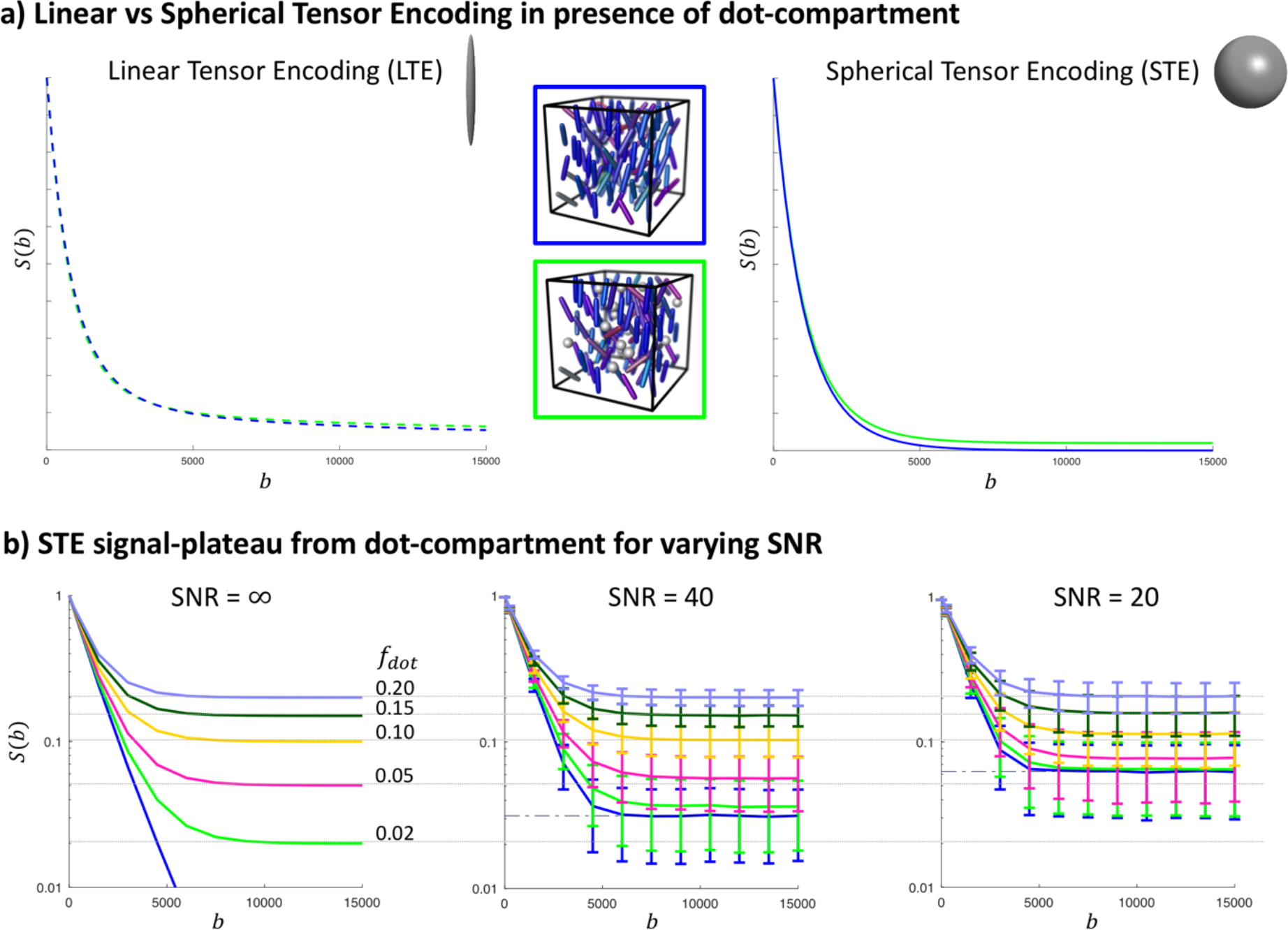
a) Simulations of LTE and STE data for two different scenarios (schematically represented in the middle): dispersed sticks representing axons surrounded by extra-axonal space (top, blue surround), *vs* dots + dispersed sticks surrounded by extra-axonal space (bottom, green surround). Here, we used a Watson distribution to simulate a stick orientation dispersion (OD) of 0.7 and *f*_*dot*_ = 0 (blue), and OD = 0.5 and *f*_*dot*_ = 0.02 (green). The two scenarios result in very similar signals for LTE across a wide b-value range, and can be disentangled better at high b-values with STE. A linear y-scale is chosen here to not make small differences seem disproportionally large. b) STE simulated as in (a) but with varying *f*_*dot*_, at different SNR levels. The dashed-dotted line represents the rectified noise floor, and the error bars represent the mean and standard deviation over 5000 noise realisations. A logarithmic y-scale is chosen here to improve the visualisation for different *f*_*dot*_. b is given in s/mm^2^.

Previous work using STE obtained by a series of pulsed gradients on a clinical system concluded that the dot-signal-fraction is likely lower than 2% in WM, and therefore has a negligible contribution to the dMRI signal (Dhital et al., 2018). However, the gradient amplitude available on clinical MRI scanners (40-80 mT/m) limits the maximal b-value per unit signal-to-noise ratio (SNR) – needed for reliable quantification of the dot-signal-fraction – whereas ultra-strong gradients (e.g., 300 mT/m) allow much higher b-values per unit SNR (Jones et al., 2018; Setsompop et al., 2013). Furthermore, the previous implementation of STE used waveforms with exceedingly low efficiency (Sjölund et al., 2015). In this work, we leverage the power of ultra-strong gradients and optimised asymmetric STE gradient waveforms to reduce the echo time (TE) significantly, thereby increasing SNR. This allows signal decays to be examined in the living human brain over a much larger range of b-values typically unachievable using clinical MRI scanners, and thus provides a more reliable assessment of signal fractions which could result from isotropically-restricted compartments. In addition, we extend the analysis to tissue types beyond cerebral WM, including deep grey matter (GM) and the cerebellum.

## 2. Theory

Assuming Gaussian diffusion within a compartment, the signal *S*_*i*_ arising from the *i*^th^ compartment, represented by diffusion tensor ***D***_*i*_ and which contributes a relative signal fraction *f*_*i*_ to the signal, probed by b-tensor ***B*** can be described by

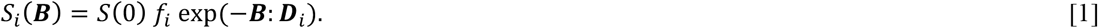

The total signal is then the sum of the signals from the individual compartments, with *f*_*i*_ summing to one. The b-tensor is a positive semi-definite tensor which we here design to be axially symmetric; it can then be characterised by its trace *b* = Tr(***B***) = (*b*_∥_ + 2*b*_⊥_) – better known as the b-value, *b*, – and its anisotropy *b*_Δ_ = (*b*_∥_ − *b*_⊥_) / (*b*_∥_ + 2*b*_⊥_) (Eriksson et al., 2013; Topgaard, 2017; Westin et al., 2016), where *b*_∥_ and *b*_⊥_ are the eigenvalues corresponding to the eigenvectors along and perpendicular to the symmetry axis, respectively. *S*(0) represents the signal at *b* = 0 s/mm^2^, and ***B***: ***D*** denotes the inner product between the tensors.

In the case of STE, the b-tensor is isotropic and thus *b*_Δ_ = 0. For *n* non-exchanging Gaussian compartments, the STE-signal simplifies to

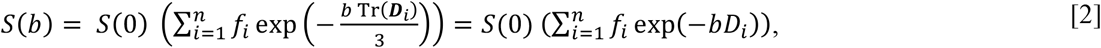

where *D*_*i*_ = Tr (***D***_*i*_) /3 is the mean apparent diffusivity of each compartment.

An isotropically restricted compartment typically exhibits a very low mean apparent diffusivity. If we index this compartment as *i* = 1 and assume *D*_1_ ≪ *D*_*i*_, *i* = 2, …, *n*, then the only remaining signal when approaching high b-values (beyond a certain b-value, *b*_*s*_) is that arising from the isotropic restricted compartment:

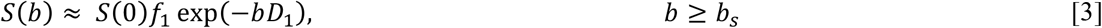

For example, for a two-compartment system with *D*_1_ = 0.1 µm^2^/ms and *D*_2_ = 0.8 µm^2^/ms, the signal from the second compartment is reduced to ~0.1 % for *b*_*s*_ = 8500 s/mm^2^, while the signal from the first compartment is only reduced to 42 %. This means that the behaviour of *S*(*b*) at increasing b-values is increasingly dominated by compartments with lower apparent diffusivity.

In the case of a dot-compartment with zero mean apparent diffusivity, i.e. *D*_*dot*_ = 0, Eq. [3] simplifies to

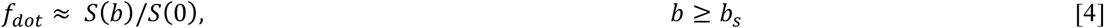

such that the dot-signal-fraction is equal to the relative signal that remains at high b-values. Fig.1b shows the simulated signal in the case of non-exchanging compartments of which one is a dot-compartment. Even if the signal does not yet exhibit a plateau, the *relative* signal at the highest b-value can serve as an upper limit of *f*_*dot*_, because *f*_*dot*_ ≤ *S*(*b*_*max*_)/*S*(0). The accuracy of this limit is affected by the presence of the rectified noise floor, 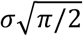, with σ standard deviation of the Gaussian noise added to each of the real and imaginary channels (Jones and Basser, 2004) (Fig. 1b).

## 3. Methods

### 3.1 Data

Five healthy adult volunteers were included in the study (3 female), which was approved by the Cardiff University School of Medicine ethics committee. Written informed consent was obtained from all participants.

Participants were scanned on a 3T Connectom MRI system (Siemens Healthcare, Erlangen, Germany) with an ultra-strong 300 mT/m gradient set. The acquisition protocol included a structural MPRAGE (Magnetization Prepared RApid Gradient Echo) (de Lange et al., 1991) and dMRI sequences. The dMRI data were acquired using a prototype spin-echo sequence with an echo-planar imaging (EPI) readout, that enables user-defined gradient waveforms to be used for diffusion encoding (Szczepankiewicz et al., 2019a). For STE we used b = [250, 1500, 3000, 4500, 6000, 7500, 9000, 10500, 12000, 13500, 15000] s/mm^2^, repeated [6, 9, 12, 15, 18, 21, 24, 27, 30, 33, 36] times, respectively. The b-values and repetitions were interleaved over volumes to reduce the impact of system drift (Hutter et al., 2018; Vos et al., 2016). For LTE, the b-tensor principal eigenvectors were distributed over the unit sphere for each b-shell. b = 0 s/mm^2^ (b0) images were acquired every 15^th^ image for monitoring and correction of subject motion. Additional b0 images with reversed phase-encoding were acquired to correct for susceptibility distortions (Chang and Fitzpatrick, 1992). No in-plane acceleration was used, and imaging parameters were: voxel size = 4×4×4 mm^3^, matrix = 64×64, 34 slices, TR = 4300 ms, partial-Fourier = 6/8, bandwidth = 1594 Hz/pixel.

The waveforms used for STE and LTE are shown in Fig. 2, and were optimised numerically (Sjölund et al., 2015) to be Maxwell-compensated (Szczepankiewicz et al., 2019b) and enable a TE as short as 88 ms. These waveforms render superior encoding efficiency due to their optimised asymmetric trajectory in q-space compared to standard 1-scan-trace imaging (which requires TE = 270 ms for b = 15000 s/mm^2^).

**Fig. 2:**
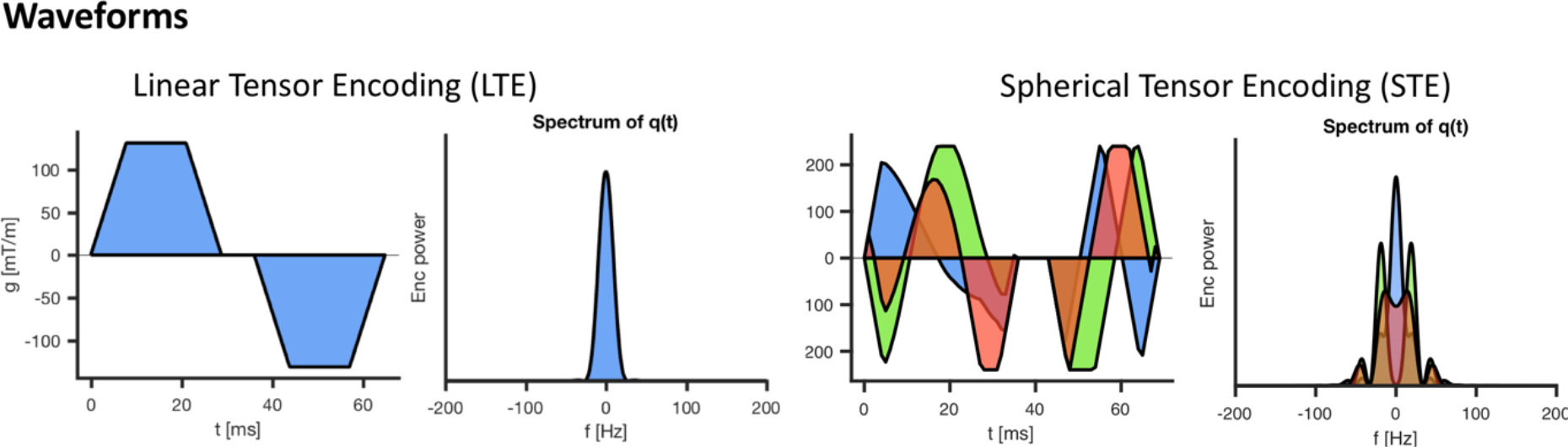
Linear tensor encoding (LTE) and spherical tensor encoding (STE) waveforms for b = 15000 s/mm^2^, and the corresponding power spectra of the dephasing vector *q*. Timings for the first waveform, temporal gap (180° pulse), and second waveform were [28.6, 6.9, 28.6] ms for LTE and [35.5, 6.9, 25.6] ms for STE. The maximum gradient amplitudes along a single axis were 131 and 240 mT/m for LTE and STE, respectively.

### 3.2 Preprocessing

The dMRI data were corrected for Rician noise bias (Koay et al., 2009a; St-Jean et al., 2016) using estimates of the Gaussian noise standard deviation (Koay et al., 2009b) and the true underlying Rician signal (Veraart et al., 2016), to determine whether or not any plateau arising in the signal decay curve could be attributed to the effects of the noise floor. The data were checked for signal intensity errors including slice-wise outliers (Sairanen et al., 2018). The STE data were corrected for subject motion by registering the interleaved b0 images to the first b0 image and applying the corresponding transformations to the diffusion-weighted images (DWIs). The LTE data were corrected for subject motion and eddy-current geometrical distortions using FSL EDDY (Andersson and Sotiropoulos, 2016). Susceptibility geometrical distortions were corrected using TOPUP (Andersson et al., 2003) and for geometrical distortions due to gradient nonlinearities using code kindly provided by colleagues at the Athinoula A. Martinos Center for Biomedical Imaging at Massachusetts General Hospital (Glasser et al., 2013; Jones et al., 2018; Rudrapatna et al., 2018; Setsompop et al., 2013).

The MPRAGE image was segmented into regions using Freesurfer (Fischl et al., 2002) and affinely-registered to the corrected b0 image using FSL FLIRT (Jenkinson et al., 2012). The resulting WM, GM, deep GM (dGM), cerebellar WM (cWM) and cerebellar GM (cGM) segmentations were then used to guide the delineation of regions-of-interest (ROIs) for further analysis. Only voxels in which the tissue probability derived from the Freesurfer segmentations was larger than 90% were considered, and the ROIs were drawn manually to avoid including signal artefacts. For WM, two separate regions were considered: ROIs were drawn on coronal slices in medial WM lateral to the midbody of the corpus callosum (denoted by mWM), and in the occipital regions (denoted by oWM), see Fig. 4.

### 3.3 Quantitative characterisation of the STE signal at high b-values

Eq. [3] was fitted to the data with *b*_*s*_ = 10000 s/mm^2^, using a nonlinear least-squares trust-region-reflective algorithm implemented in MATLAB (The MathWorks, Natick, USA). The fit was randomly initialised 10 times, and the solution with the lowest residual norm was selected. In addition, estimates 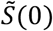 and 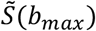 were obtained, from which 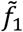, and an upper limit 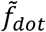 were derived according to Eqs. [3] and [4], respectively.

## 4. Results

### 4.1 ROI delineation

Fig. 3 shows the Freesurfer segmentation results overlaid on individual diffusion-weighted images of one participant. Fig. 4 shows results of the manually delineated ROIs visualised for one of the healthy subjects. The mWM, oWM, cWM, cGM, and dGM ROIs include on average 59, 58, 39, 199, and 52 voxels across participants, respectively. The GM segmentations only include a few voxels that are classified as > 90% GM, which are sparsely distributed. We therefore only consider data in the mWM, oWM, cWM, cGM, and dGM ROIs.

**Fig. 3:**
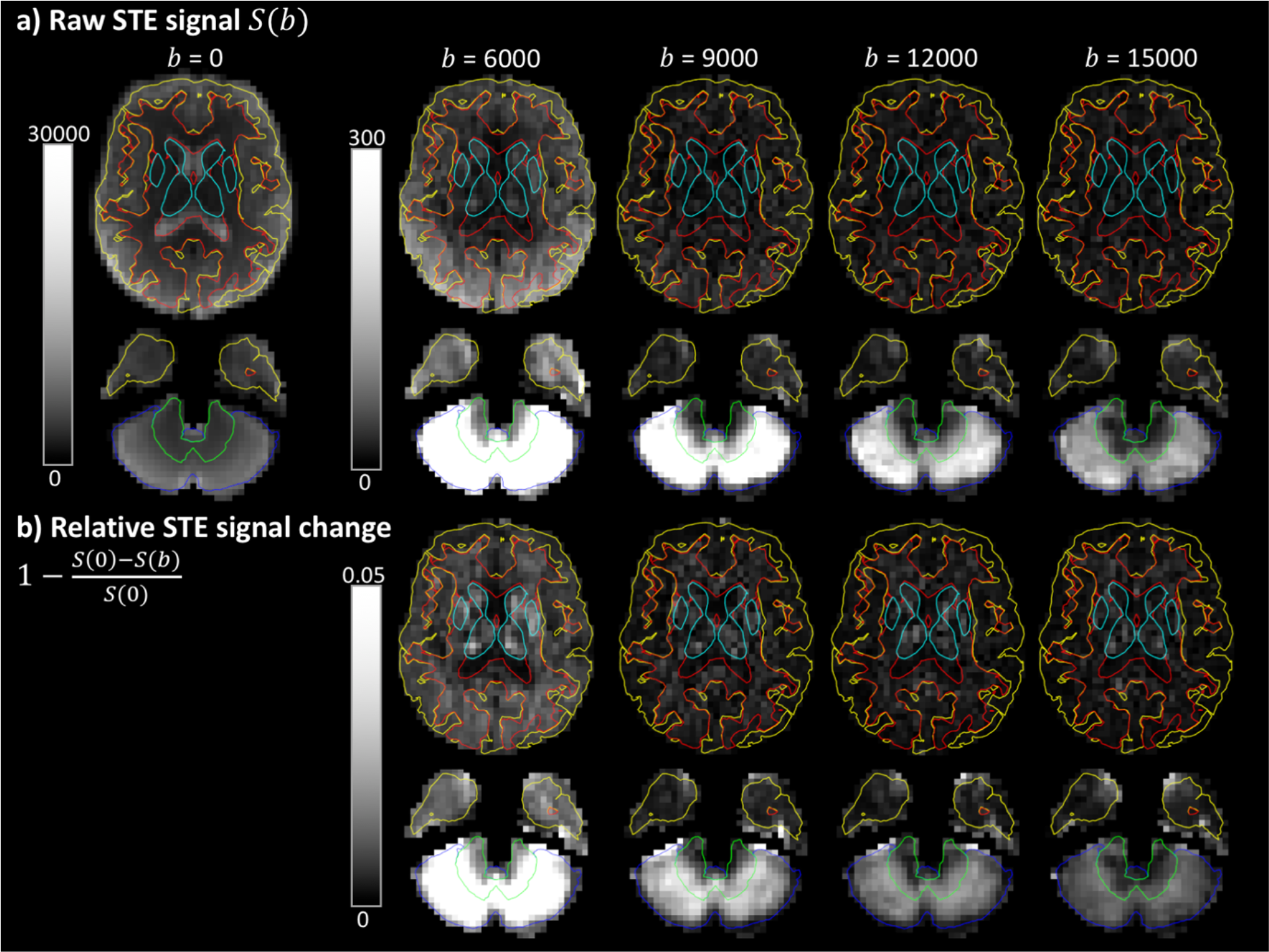
a) STE signal as a function of b-value (in s/mm^2^), with the Freesurfer tissue segmentations indicated in red = WM, yellow = GM, cyan = deep GM, green = cerebellar WM and blue = cerebellar GM. b) Relative STE signal change.

**Fig. 4:**
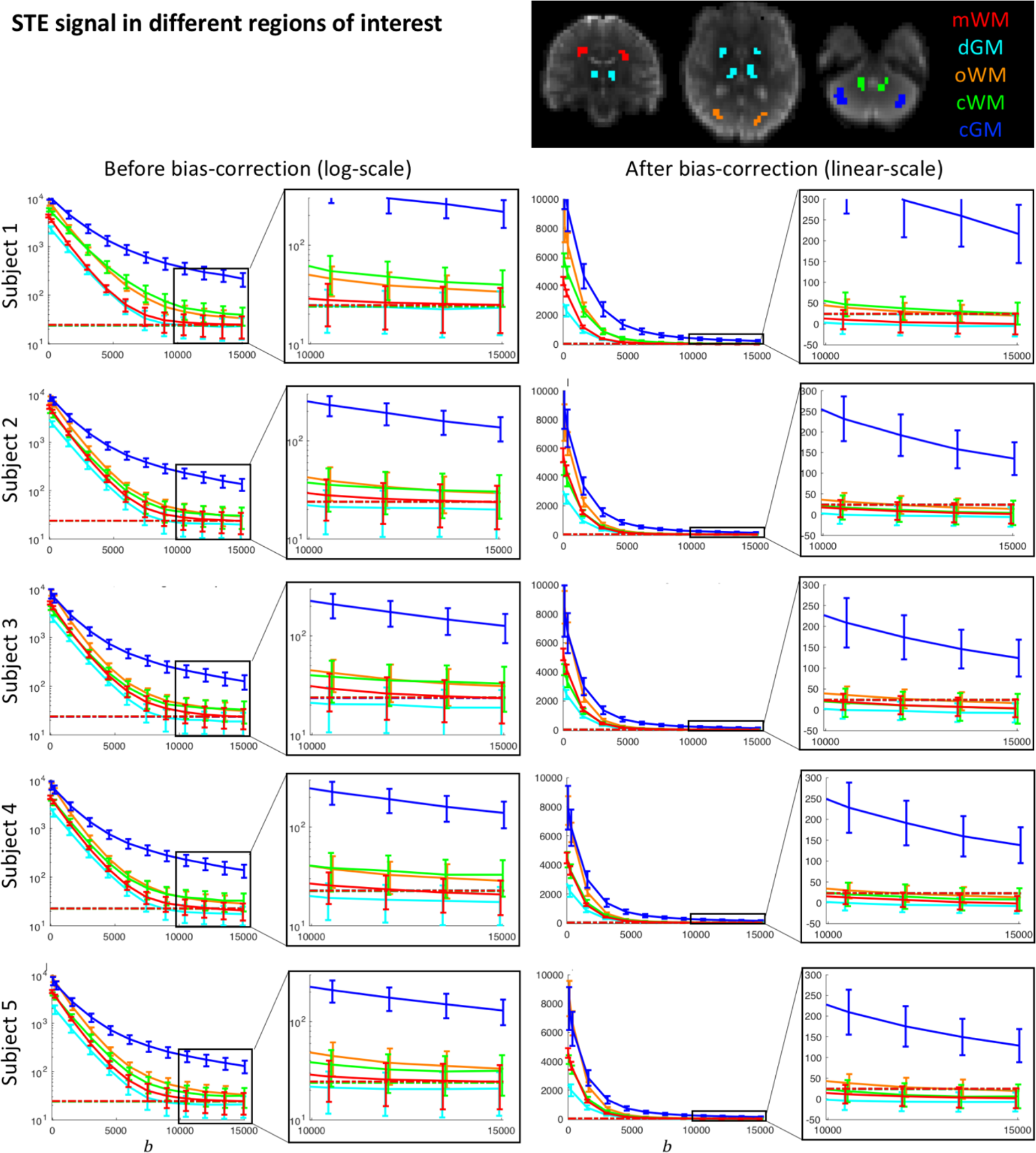
STE signal decay for 5 healthy subjects, in ROIS in the mWM (red), oWM (orange), cWM (green), dGM (cyan), and cGM (blue); examples of the ROIs are shown for Subject 1. The left column shows the signal before Rician-bias correction plotted with a logarithmic y-scale, to better visualise deviations from mono-exponential behaviour, with a close-up at high b-values. The right column shows the signal after Rician-bias correction plotted with a linear y-scale to be able to visualise negative values. b is given in s/mm^2^.

### 4.2 STE signal decay across all b-values

Fig. 3a shows the signal of the image intensity in individual DWIs as a function of b-value in a healthy brain. The signal intensity in most of the cerebral WM has decayed substantially at b > 10000 s/mm^2^. However, the cerebellar GM retained a remarkably high signal at these high b-values, remaining well above the noise floor even at b = 15000 s/mm^2^. Fig. 3b shows the signal in a more quantitative fashion; regions with lower intensity have a higher relative signal change compared to the *S*(0) signal. The cerebellar GM persistently has a high intensity compared to other regions and thus the lowest relative signal change.

Fig. 4 shows the signal decay for each ROI in the five healthy subjects. Both the original and Rician-bias-corrected signal decay curves are shown, accompanied by an estimate of the noise floor. At b > 10000 s/mm^2^, the signals from mWM and dGM clearly approach the noise floor. In contrast, oWM, cWM, and cGM exhibit a mean-signal that is above the noise floor for all five subjects. After Rician-noise-bias-correction, the signal is still above zero albeit it can be seen that it continues to decay.

### 4.3 STE signal characterisation at high b-values

Table 1 gives quantitative features related to the STE signal decay at high b-values. For each parameter, the median and 10^th^ – 90^th^ percentiles are given. The third column presents estimates of the relative rectified noise floor, derived from estimates of the noise standard deviation and b0 signal (i.e. 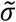 and 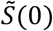). An estimate of the relative noise floor of 0.5% indicates that an SNR > 250 on the b0 signal could be achieved. The fourth column presents estimates for the dot-signal-fraction 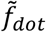(Eq. [3]) and the last two columns present estimates in the case of an isotropically-restricted compartment with non-zero diffusivity (Eq. [4], i.e. 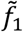, and 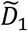). All estimates are obtained after Rician bias-correction to reduce bias from the least-squares fitting. We will describe characteristics of these features for the different ROIs in the following paragraphs.

**Table 1:**
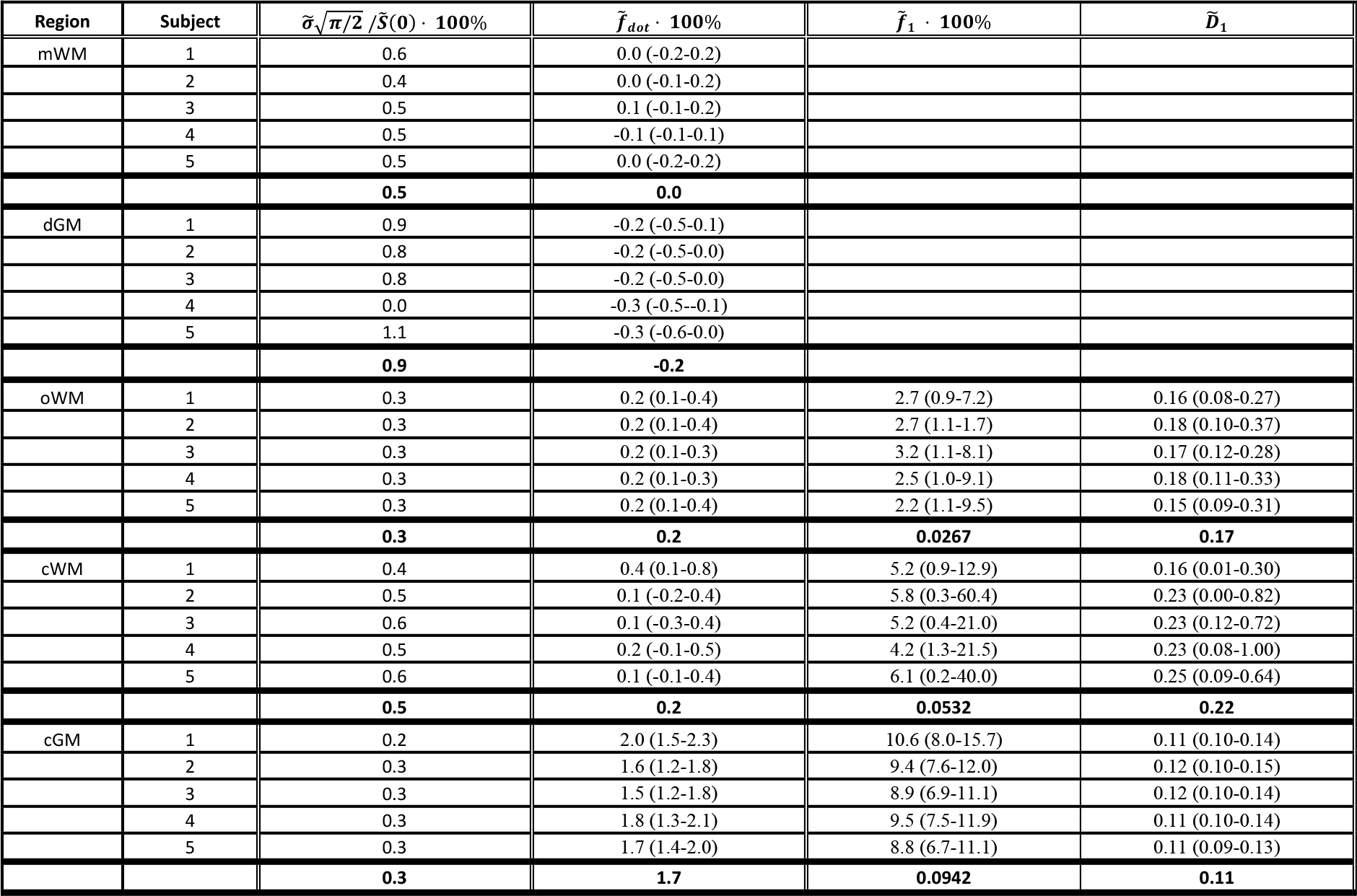
Parameter estimates (median and 10-90 percentile) for the standard deviation 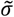,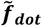,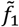 and 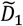, in different ROIs. *D* has units of µm^2^/*ms*.

| Region | Subject | $\tilde{\sigma} \cdot \sqrt{\pi/2} / \tilde{S}(0) \cdot 100\%$ | $\tilde{f}_{dot} \cdot 100\%$ | $\tilde{f}_1 \cdot 100\%$ | $\tilde{D}_1$ |
| --- | --- | --- | --- | --- | --- |
| mWM | 1 | 0.6 | 0.0 (-0.2-0.2) |  |  |
|  | 2 | 0.4 | 0.0 (-0.1-0.2) |  |  |
|  | 3 | 0.5 | 0.1 (-0.1-0.2) |  |  |
|  | 4 | 0.5 | -0.1 (-0.1-0.1) |  |  |
|  | 5 | 0.5 | 0.0 (-0.2-0.2) |  |  |
|  |  | **0.5** | **0.0** |  |  |
| dGM | 1 | 0.9 | -0.2 (-0.5-0.1) |  |  |
|  | 2 | 0.8 | -0.2 (-0.5-0.0) |  |  |
|  | 3 | 0.8 | -0.2 (-0.5-0.0) |  |  |
|  | 4 | 0.0 | -0.3 (-0.5--0.1) |  |  |
|  | 5 | 1.1 | -0.3 (-0.6-0.0) |  |  |
|  |  | **0.9** | **-0.2** |  |  |
| oWM | 1 | 0.3 | 0.2 (0.1-0.4) | 2.7 (0.9-7.2) | 0.16 (0.08-0.27) |
|  | 2 | 0.3 | 0.2 (0.1-0.4) | 2.7 (1.1-1.7) | 0.18 (0.10-0.37) |
|  | 3 | 0.3 | 0.2 (0.1-0.3) | 3.2 (1.1-8.1) | 0.17 (0.12-0.28) |
|  | 4 | 0.3 | 0.2 (0.1-0.3) | 2.5 (1.0-9.1) | 0.18 (0.11-0.33) |
|  | 5 | 0.3 | 0.2 (0.1-0.4) | 2.2 (1.1-9.5) | 0.15 (0.09-0.31) |
|  |  | **0.3** | **0.2** | **0.0267** | **0.17** |
| cWM | 1 | 0.4 | 0.4 (0.1-0.8) | 5.2 (0.9-12.9) | 0.16 (0.01-0.30) |
|  | 2 | 0.5 | 0.1 (-0.2-0.4) | 5.8 (0.3-60.4) | 0.23 (0.00-0.82) |
|  | 3 | 0.6 | 0.1 (-0.3-0.4) | 5.2 (0.4-21.0) | 0.23 (0.12-0.72) |
|  | 4 | 0.5 | 0.2 (-0.1-0.5) | 4.2 (1.3-21.5) | 0.23 (0.08-1.00) |
|  | 5 | 0.6 | 0.1 (-0.1-0.4) | 6.1 (0.2-40.0) | 0.25 (0.09-0.64) |
|  |  | **0.5** | **0.2** | **0.0532** | **0.22** |
| cGM | 1 | 0.2 | 2.0 (1.5-2.3) | 10.6 (8.0-15.7) | 0.11 (0.10-0.14) |
|  | 2 | 0.3 | 1.6 (1.2-1.8) | 9.4 (7.6-12.0) | 0.12 (0.10-0.15) |
|  | 3 | 0.3 | 1.5 (1.2-1.8) | 8.9 (6.9-11.1) | 0.12 (0.10-0.14) |
|  | 4 | 0.3 | 1.8 (1.3-2.1) | 9.5 (7.5-11.9) | 0.11 (0.10-0.14) |
|  | 5 | 0.3 | 1.7 (1.4-2.0) | 8.8 (6.7-11.1) | 0.11 (0.09-0.13) |
|  |  | **0.3** | **1.7** | **0.0942** | **0.11** |

For the mWM and dGM ROI, the mean signal at high b-values converges to the noise floor (Fig. 4). We estimate an upper limit of *f*_*dot*_ of 0.5% and 0.9% respectively.

For the oWM and cWM ROI, the estimated upper limits of *f*_*dot*_ are 0.3% and 0.5% respectively. The signal at high b-values is still decaying, and can thus be better explained by the presence of a compartment with non-zero apparent diffusivity with estimated signal fractions of 2.7% and 5.3%, and estimated apparent mean diffusivities of 0.17 and 0.22 µm^2^/ms for oWM and cWM, respectively. Fig. 5a shows scatter plots of these estimates, showing that the spread is large (see also the percentiles in Table 1).

**Fig. 5:**
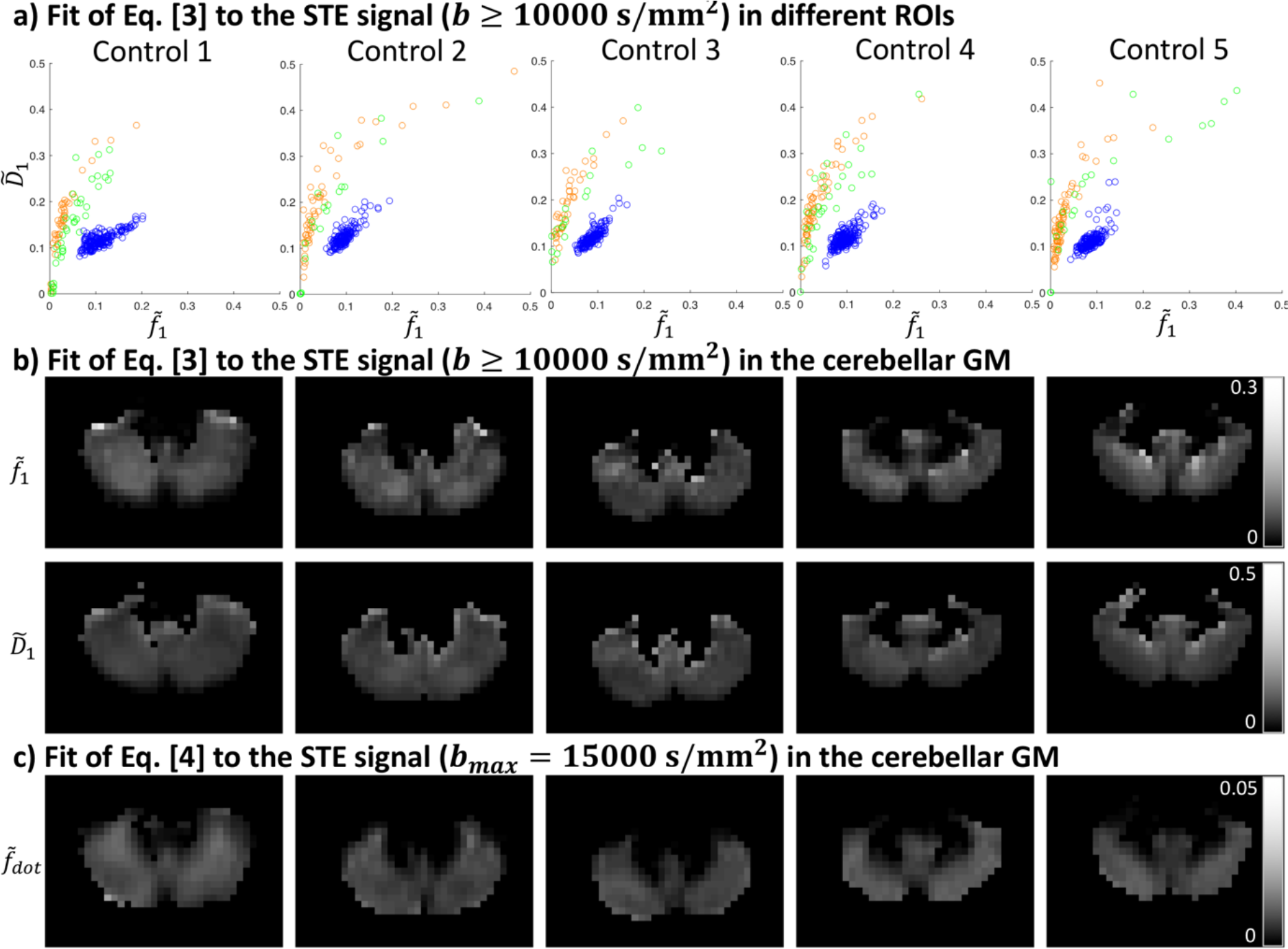
a) Parameter estimates 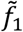 and 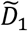, in the oWM (orange), cWM (green), and cGM (blue) ROIs. b-c) Map of the fits of Eqs. [3] (not assuming zero apparent diffusivity) and [4] (assuming zero apparent diffusivity) in an axial slice of the cGM, respectively; the cerebellar WM is masked out. *D* has units µm^2^/ms.

For the cGM ROI, we find an upper limit of *f*_*dot*_ of 1.7%, with a residual signal that is well above the noise floor. Per Eq. [3], we estimate an average apparent mean diffusivity of 0.11 µm^2^/ms and an average signal fraction of 9.4%. In some areas, the signal fraction is estimated as high as 16%. These estimates are consistent across healthy subjects (Fig. 5a). When visualising the estimates in the cerebellar GM one can observe that they are spatially heterogeneous (Fig. 5b). As a comparison, we show the spatial variability of *f*_*dot*_ in Fig. 5c.

### 4.4 Comparison of LTE and STE signals

In Fig. 6 one can readily appreciate the difference between b0-normalised STE and directionally-averaged LTE signals in the different tissue types. These diffusion weightings also give complementary information in GM, where the STE encoding at high b-values has suppressed signal arising from compartments that are mobile along at least one axis (e.g., ‘sticks’ that could represent axons). The overlap of the signal decay curves is high between the healthy controls.

**Fig. 6:**
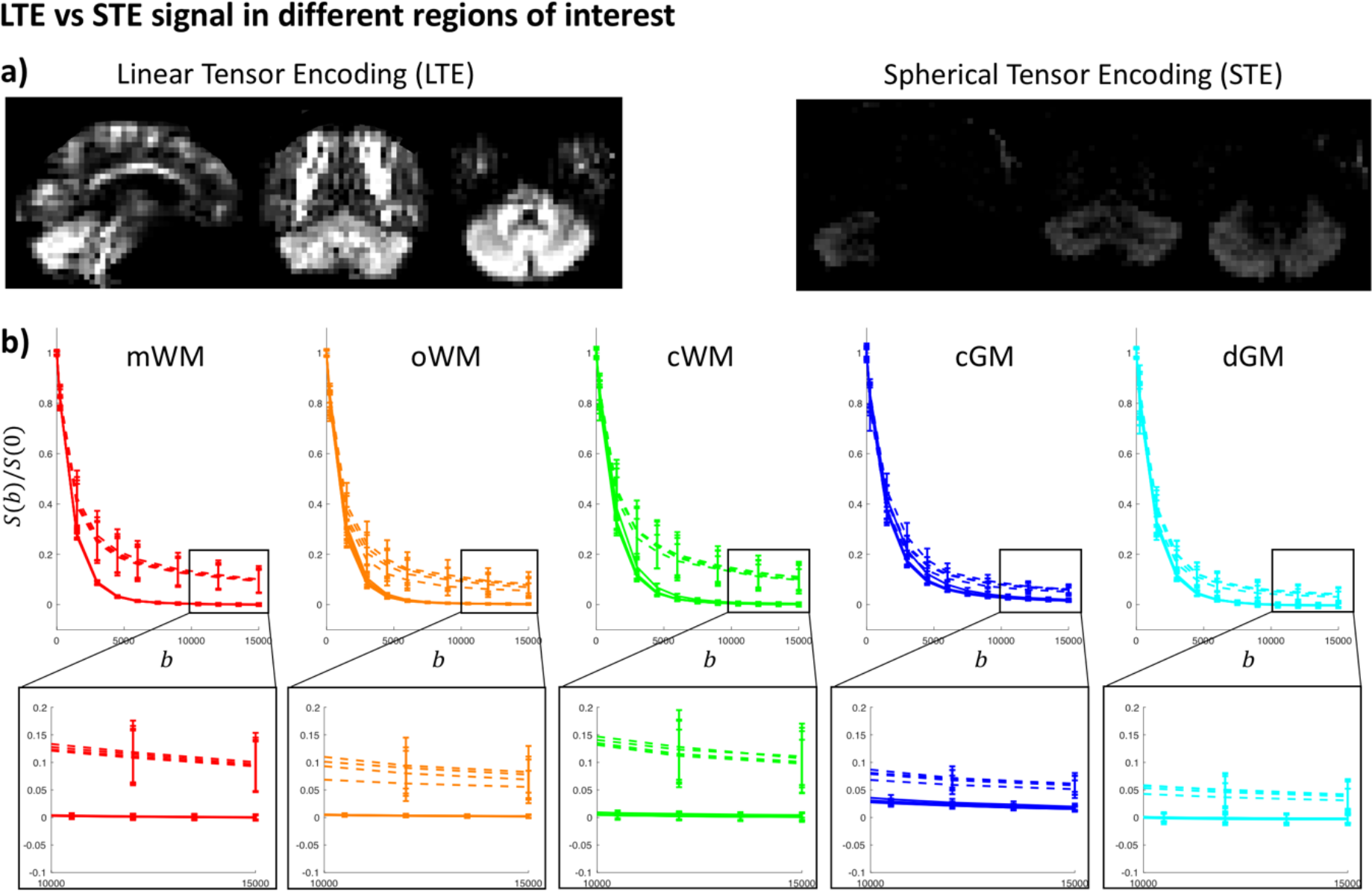
a) Signal upon LTE and STE (b = 15000 s/mm^2^) with the same intensity scale. b) LTE (dashed lines) and STE (solid lines) signals, with b in s/mm^2^. Colours correspond to Fig. 4.

## 5. Discussion

In this study we report for the first time a clear depiction *in vivo* from an isotropically-restricted compartment in dMRI. This compartment is present particularly in the cerebellar GM, but support for its existence can also be found in the WM. Our observations were enabled by ultra-strong gradient hardware (Jones et al., 2018; Setsompop et al., 2013) and recent developments for tensor-valued diffusion encoding (Sjölund et al., 2015; Szczepankiewicz et al., 2019b). STE provides essential complementary information to LTE, but the waveforms generally take up more time than Stejskal-Tanner LTE encoding, leading to long TEs and thereby inferior SNR. With the help of ultra-strong diffusion gradients (240 mT/m along a single axis, Fig. 2), a TE as short as 88 ms could be achieved even for a b-value of 15000 s/mm2. As a result, the SNR was above 250 in the b0 images, and we could clearly observe signal amplitudes well above the noise floor. Before going into the implications, we will first discuss some general observations.

A plateau of the diffusion-weighted signal (i.e., region of no further signal decay with increasing b-value), even at high b-values, was not observed in any region of interest. This makes the significant contribution of water residing in a dot-compartment with zero apparent diffusivity and no exchange unlikely. This observation is in agreement with previous work (Dhital et al., 2018; Veraart et al., 2019). Nevertheless, a *slowly decaying* STE signal was observed in some regions, and this observation can be supported by two hypotheses: (i) a zero-apparent-diffusivity compartment exists but is not observed as such because it is in exchange with its surroundings; or (ii) the compartment exhibits a low but non-zero apparent diffusivity. The first hypothesis is analysed in Supplementary Fig. 1, which shows the noiseless signal decay for different exchange times using a two-compartment Kärger model (Kärger, 1971; Nilsson et al., 2010). At infinite exchange times, the estimated dot-signal fraction approaches its true value. However, at exchange times of e.g. 500 ms the signal does not exhibit a plateau and the estimated upper limit of *f*_*dot*_ is negatively biased.

Regarding the second hypothesis, a slow-diffusing component has not been observed previously in STE data. Previous work has characterised mean apparent diffusivities derived from STE data up to ~b = 6000 s/mm^2^ by using a regularised inverse Laplace transform (Avram et al., 2019) or by fitting a finite series of exponentials that could represent different compartments and comparing the fits of the models through the Akaike Information Criterion (AIC) (Dhital et al., 2018). These works showed little deviation from mono-exponential behaviour in WM and single-peak diffusivity distributions in brain parenchyma in the range of b-values used. However, in the logarithmic plots in Fig. 4 one can clearly observe that the signal decay starts deviating from mono-exponential behaviour for ~b > 5000 s/mm^2^ in most tissue types, which could explain why this component has not been reported previously.

### 5.1 Signal representation and implications

Rather than quantifying the signal across the entire range of b-values and comparing the fit of models with different numbers of compartments, we focus here on quantifying the STE signal at high b-values using a simple representation based on the often-adopted assumptions of Gaussian diffusion and no exchange. Under these assumptions, the results provide support for the presence of an isotropic water pool with low diffusivity in the oWM, cWM, and cGM ROIs. In WM, Dhital et al. (2018) found that for a hypothetically small, yet finite, diffusivity of *D*_1_ = 0.1 µm^2^/ms, the estimated dot signal fraction was 2.7%. In the present study, we found a similarly low signal fraction, but the diffusivity was estimated to be twice as high (~0.2 µm^2^/ms) in the oWM and cWM ROIs, albeit with a high uncertainty (Table 1). In the medial WM the signal converged to the noise floor; this could be caused by the larger distance to the RF receiving coils (and thus lower SNR), or a genuinely lower density of slow-diffusing components compared with the occipital WM, or both.

In the cGM ROI, the signal fraction of the slowly diffusing isotropic water pool was estimated to be as large as 16%, and this component thus makes a significant contribution to the signal. Linking this finding to tissue microstructure derived from histology or realistic numerical simulations of brain cells (Palombo et al., 2019) is the subject of future work. It has been suggested previously that in cortical GM, the abundance of cell bodies has a significant impact on the LTE signal at high b-values (Palombo et al., 2018). In that work, the LTE signal at high b-values was considered to be arising from non-exchanging sticks representing neurites, and spheres with a finite radius representing cell bodies. Following this picture, STE at high b-values would nullify the stick-signal and only the signal specific to the cell bodies would remain.

The cerebellum has an important role in motor coordination, but it is becoming increasingly apparent that it also has an active role in cognition and emotion (O’Halloran et al., 2012; Tedesco et al., 2011). The neurons in the cerebellar cortex are highly organised, consisting of densely-packed granule cells and larger Purkinje cells with a cloud of dendritic spines. One can speculate that the isotropically-restricted signal comes from within small spaces such as the granule cells or dendritic spines. While this hypothesis remains to be validated, studying the STE signal provides exciting avenues for gaining further insight into changes in tissue microstructure in disorders associated with the cerebellum. dMRI studies have already shown changes in ataxia (Dayan et al., 2016; Salvatore et al., 2014), Parkinson’s disease, and Alzheimer’s disease (Mormina et al., 2017), where metrics such as mean diffusivity and diffusion-tensor (DT)-derived fractional anisotropy (FA) were studied. These studies mostly focused on cerebellar WM (e.g. peduncles). Recently, measures beyond the DT have been derived in cerebellar WM and GM, with the aim of being more specific to different compartments and the underlying neurobiology (Savini et al., 2018). Fig. 5b shows spatial variability in the estimated parameter maps. In future work we aim to look at the variability across and within different lobules, by registration to atlases (Diedrichsen et al., 2009).

The use of pulsed-gradients allows a more precise definition of the time-scale of diffusion. The STE waveforms in Fig. 2 have broader frequency spectra, affecting the way time-dependent diffusion is encoded (Lundell et al., 2018). Under the assumption of Gaussian (and thus time-independent) diffusion in each compartment (as in Section 2), the net signal becomes non-monoexponential but remains time-independent; as such the signal decay arising from two sets of waveforms with the same B-tensor, but different frequency spectra, would look identical. However, the assumption of compartmental Gaussian diffusion is theoretically only valid for sufficiently short or long diffusion times or low diffusion weightings; beyond these regimes time-dependent diffusion will be encoded differently by waveforms with different frequency spectra. The use of different LTE waveforms with different frequency characteristics and b-values up to 5000 s/mm^2^ has previously revealed a strong contrast in the cerebellum (Lundell et al., 2017, 2015). In the specific case of STE as studied here, several works (de Swiet and Mitra, 1996; Jespersen et al., 2019; Lundell et al., 2018) have shown that non-Gaussian diffusion within each compartment can lead to anisotropic time-dependence, i.e., probing different time-dependence in different directions. This means that for anisotropic pores, such as cylinders and ellipsoids, the signal decay in STE still depends on the orientation and dispersion of the pores (or the rotation of the waveforms). In the present study, we focused on the high b-value regime to suppress the signal from anisotropic compartments that have significant mobility along at least one axis. Therefore, the remaining signal is expected to come from restricted isotropic compartments only, and is as such expected to be rotationally invariant. Non-Gaussian diffusion within these isotropic compartments becomes a contributing factor if one for example tries to estimate the variance of the highly restricted isotropic diffusivities (Jespersen et al., 2019), which is beyond the aim of this study.

### 5.2 SNR and spatial resolution

The spatial resolution used here to achieve the necessary SNR (i.e. voxel size 4×4×4 mm^3^) is relatively coarse, especially if one tries to study highly curved structures such as the cortical grey matter. Super-resolution and gSlider-SMS (Setsompop et al., 2018) diffusion acquisitions provide exciting future avenues for increasing the spatial resolution while maintaining sufficient SNR. Compared with LTE super-resolution reconstruction, which has been extended recently to incorporate the angular relation between different diffusion measurements (Van Steenkiste et al., 2016), STE super-resolution would theoretically be more straightforward as the need to vary the orientation of the principal eigenvectors of the B-tensor is obviated.

### 5.3 Pre-processing

The low SNR of the STE data at high b-values made the pre-processing of the data challenging. dMRI pre-processing pipelines typically include motion correction and geometric distortion correction. The geometric distortions generally include those resulting from eddy currents and susceptibility differences, and the use of strong gradients requires an additional step to correct for any possible geometric distortions arising from gradient nonlinearities. Subject motion and eddy-current geometric distortions in high b-value data are often corrected for using a prediction-based framework (Andersson et al., 2017; Ben-Amitay et al., 2012); high b-value images are predicted from the corrected low b-value images, and the acquired high b-value images are subsequently registered to the predicted images. Strategies to predict high b-value data with different B-tensors from low b-value data are available (Nilsson et al., 2015), but the deformations allowed at high b-values have to be fairly constrained because only a relatively low signal can be observed in only few regions. When applying tools optimised for LTE images and/or moderate b-value STE images, we observed suspiciously large deformations in the high b-value STE data that could not be verified. In this study, we therefore opted for a conservative strategy where we acquired interleaved b0 images (every 15^th^ image) to correct for subject motion in STE data. This necessarily led to differences in the processing of LTE and STE data; i.e., the STE data were only corrected with a rigid transformation which cannot account for higher order deformations e.g., due to eddy currents. While, theoretically, the eddy current deformations between STE images of the same b-value should be similar, future work should be attributed to optimising the processing of high b-value STE data. Future work will furthermore focus on collecting complementary information by means of real-time motion tracking (e.g. optical tracking (Qin et al., 2009)) and dynamic field measurements (De Zanche et al., 2008) to provide robust correction for subject motion and geometrical distortions in these data.

In this work we corrected for geometric distortions arising from gradient nonlinearities, but gradient nonlinearities additionally cause spatiotemporally varying B-tensors. Strategies have been developed to take this into consideration, which were mostly evaluated on data acquired with Stejskal-Tanner encoding (Bammer et al., 2003; Glasser et al., 2013; Jones et al., 2018; Rudrapatna et al., 2018). Future work will be attributed to investigating the effect of gradient nonlinearities on the signal arising from free waveforms, as used in this study.

Correcting for the Rician noise bias is of importance here to obtain accurate estimates of the parameters in Eqs. [3-4] when using least-squares optimisation. Here we used the approach of Koay et al. (2009a) which relies only on magnitude data. When phase data are available, this can alternatively be leveraged to obtain Gaussian-distributed data (Eichner et al., 2015; Pizzolato et al., 2016). The process of Rician debiasing can yield signal estimates below the noise floor. To evaluate the accuracy and precision of this approach, we applied the same debiasing step as described in Section 3.2 to the simulated data of Fig. 1 (using the same acquisition protocol as in the *in vivo* data). Supplementary Fig. 2 shows estimates of *f*_*dot*_ for different SNR, before and after Rician debiasing. Indeed, estimation before Rician debiasing results in overestimation of *f*_*dot*_. The expectation value of the error term in the case of nonlinear least squares and Rician-distributed data has been shown to converge to zero relatively slowly as a function of SNR (Veraart et al., 2013), which means that estimates can still be biased even if the SNR is larger than 2. After debiasing, our simulations indicate that signal estimates below the noise floor likely have a negative bias, resulting in a slight underestimation of *f*_*dot*_. This bias, however, converges to zero faster than prior to Rician debiasing.

## 6. Conclusion

In this work, we combined ultra-strong gradients and efficient spherical tensor encoding to study the isotropic dMRI signal at ultra-high b-values, targeting the dot-compartment. Ultra-strong gradients allowed us to significantly reduce the TE, and therefore increase SNR, when acquiring data at high b-values. We further optimised encoding efficiency and TE by using asymmetric gradient waveforms instead of pulsed-gradients. A dot-compartment with zero diffusivity and no exchange would result in the signal plateauing for sufficiently high b-values; however, we found a signal significantly deviating from zero, yet still decaying across different WM regions and in the cerebellar GM. This observation is not in line with a <u>completely</u> still and non-exchanging dot-compartment. We further studied the apparent diffusivity and signal fraction in the cerebellar GM assuming Gaussian diffusion and no exchange, finding these to be remarkably consistent across healthy controls. Future work will investigate the link between this hypothesised compartment and tissue microstructure, and investigate its potential as a biomarker in pathology affecting the cerebellar GM.

## Acknowledgements

We thank Siemens Healthineers for access to the pulse sequence programming environment, and Fabrizio Fasano from Siemens Healthineers for support. We are grateful to Umesh Rudrapatna for technical support and feedback, Samuel St-Jean and Cheng Guan Koay for useful discussions on Rician-debiasing and sharing their code, and Marco Palombo for useful discussions on cell body imaging. We thank João De Almeida Martins and Daniel Topgaard for their contributions to Fig. 1, and Sila Genc and Emre Kopanoglu for feedback on the manuscript. The work was supported by a Wellcome Trust Investigator Award (096646/Z/11/Z) and a Wellcome Trust Strategic Award (104943/Z/14/Z). The data were acquired at the UK National Facility for In Vivo MR Imaging of Human Tissue Microstructure funded by the EPSRC (grant EP/M029778/1), and The Wolfson Foundation. CMWT is supported by a Rubicon grant (680-50-1527) from the Netherlands Organisation for Scientific Research (NWO). MN is supported by the Swedish Research Council (grant no. 2016-03443), and Random Walk Imaging AB (grant no. MN15).

## Supplementary Figures

**Supplementary Fig. 1:**
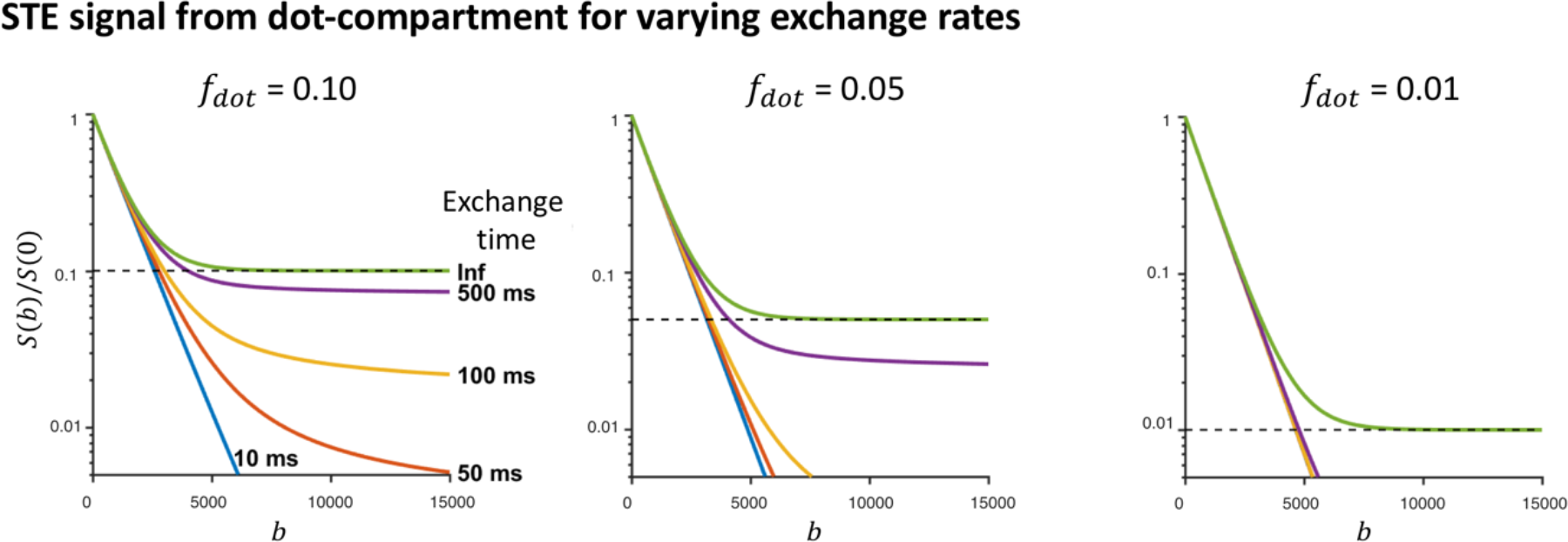
Simulated signal at variable *f*_*dot*_ and exchange times assuming a two compartment Kärger-model. Here, diffusivities were set to 1 and 0 µm^2^/ms.

**Supplementary Fig. 2:**
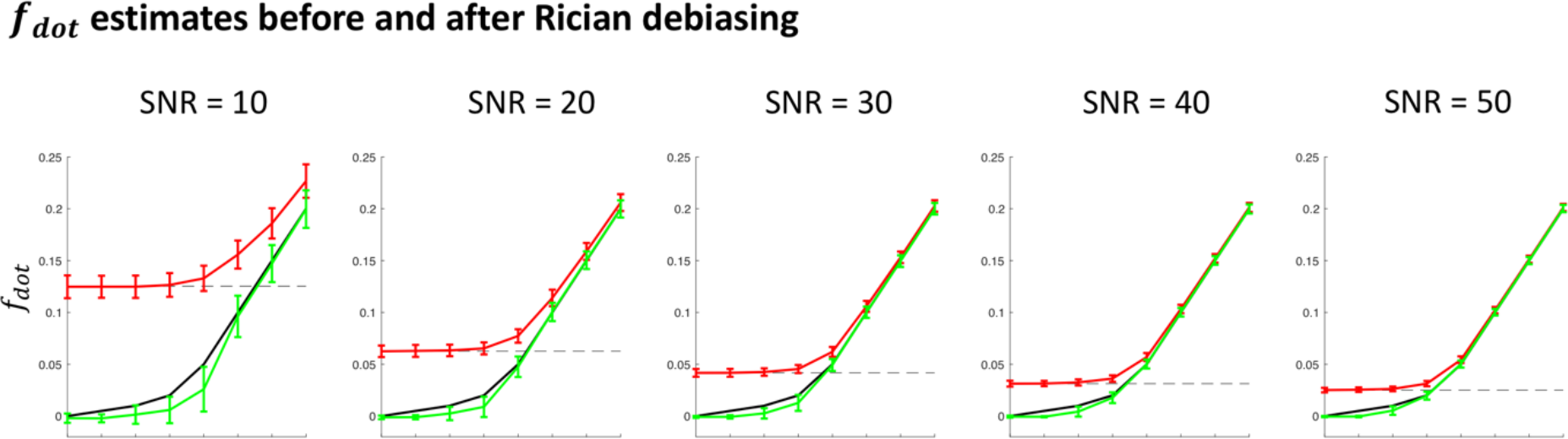
Simulated signal with different dot signal fractions (equidistantly spaced along the x-axis and connected by a line for improved visualisation, the true simulated dot signal fraction is represented by the black line and can be read from the y-axis) and different SNRs. Mean and standard deviation of the Rician signal and debiased signal are plotted in red and green respectively.

